# Fractional Hitting Sets for Efficient and Lightweight Genomic Data Sketching

**DOI:** 10.1101/2023.06.21.545875

**Authors:** Timothé Rouzé, Igor Martayan, Camille Marchet, Antoine Limasset

## Abstract

The exponential increase in publicly available sequencing data and genomic resources necessitates the development of highly efficient methods for data processing and analysis. Locality-sensitive hashing techniques have successfully transformed large datasets into smaller, more manageable sketches while maintaining comparability using metrics such as Jaccard and containment indices. However, fixed-size sketches encounter difficulties when applied to divergent datasets.

Scalable sketching methods, such as Sourmash, provide valuable solutions but still lack resource-efficient, tailored indexing. Our objective is to create lighter sketches with comparable results while enhancing efficiency. We introduce the concept of Fractional Hitting Sets, a generalization of Universal Hitting Sets, which uniformly cover a specified fraction of the *k*-mer space. In theory and practice, we demonstrate the feasibility of achieving such coverage with simple but highly efficient schemes.

By encoding the covered *k*-mers as super-*k*-mers, we provide a space-efficient exact representation that also enables optimized comparisons. Our novel tool, SuperSampler, implements this scheme, and experimental results with real bacterial collections closely match our theoretical findings.

In comparison to Sourmash, SuperSampler achieves similar outcomes while utilizing an order of magnitude less space and memory and operating several times faster. This highlights the potential of our approach in addressing the challenges presented by the ever-expanding landscape of genomic data.

SuperSampler is an open-source software and can be accessed at github.com/TimRouze/supersampler. The data required to reproduce the results presented in this manuscript is available at github.com/TimRouze/Expe_SPSP.

## 1 Introduction

The field of genomics has exploded in recent years, driven by the availability of cheap and easy sequencing data generation. The Sequence Read Archive (SRA) is a vast and to some extent under-exploited goldmine of genomic data, containing an enormous amount of genetic information. However, one of the biggest challenges in utilizing this data is the lack of efficient indexing and querying tools. The GenBank database, for example, already contains 1.2 million bacterial genomes, totaling over 5 Terabases of data. In the face of such vast genetic information, a crucial need is to promptly and precisely determine the most similar (or contained) known entry for a given query document (assembled or unassembled reads). Specifically, in this work we focus on the metagenomic assessment problem, which entails characterizing microbial communities in a specific environment using DNA sequencing data and potentially large amounts of reference entries. The complexity and diversity of the data, which contains sequences from multiple genomes, presents significant challenges.

In the metagenomic context, traditional alignment-based methods such as BLAST are increasingly computationally prohibitive due to the sheer number of potential targets for metagenome mapping. A spectrum of alignment-free techniques based on *k*-mer content have emerged as a viable alternative, with different tradeoffs. On one side of the spectrum, exact *k*-mer indexing offers linear query time [9, 14, 21], but may be too memory-intensive for large-scale applications. Probabilistic structures, when applied to large queries (on the order of kilobases), enable more scalable indexing at the expense of a random false positive rate [15, 5].

Adequate query length allows for handling false positive noise since it remains significantly lower than the required matching signal (e.g., 70% of queried *k*-mers present in the document). For extensive queries at the Megabase level, the signal strength is sufficiently robust, eliminating the need to consider all *k*-mers and enabling sublinear query time. On the other side of the spectrum, fixed-size sketches like Minhash [6], Hyperloglog [16], and Hyperminhash [29] have been effectively used for large-scale collection comparison [18, 2, 3, 30, 1]. However, they are ill-suited for divergent documents in terms of content or size, a critical limitation considering metagenomic samples typically comprise many organisms with amount of distinct *k*-mers varying by orders of magnitude.

Scaled sketches, whose size scales linearly with input size, have demonstrated better resilience to such issues. Sourmash [23], which implements scaledminhash [23] and Fracminhash [10], efficiently approximates containment and Jaccard indexes even for documents with size disparities spanning several orders of magnitude. Sourmash’s simplicity is one of its key strengths: it stores each uniformly selected *k*-mer as a 32-bit integer and compares them using a dictionary. An observation is that this technique is generic and can be applied to any type of data. Therefore, computational and memory requirements could benefit from customized selection techniques and index structures.

To this end, we propose capitalizing on the ability to represent overlapping *k*-mers with a low number of bits per *k*-mer using a *Spectrum Preserving String Set* [25]. The challenge we face is optimizing the overlap of chosen *k*-mers to achieve maximum space efficiency. To address this, we build upon the concept of super-*k*-mers [12], which are sequences of *k*-mers sharing a common selected *m*-mer called minimizer [26]. Universal Hitting Sets (UHS [19]) methods aim to design optimized *m*-mer selection schemes that cover all *k*-mers while minimizing the density of selected positions. However, our application does not require complete coverage of the *k*-mer space. Therefore, we introduce *Fractional Hitting Sets* that encompass a uniformly selected fraction of the *k*-mer space. We conduct a study on the achievable density in relation to the selected fraction and present a straightforward minimizer selection scheme that closely approaches optimal bounds. We implement this scheme in a tool called SuperSampler. The storage of enhanced super-*k*-mer sequences, partitioned by minimizers, facilitates space and time-efficient *k*-mer set comparisons. Our evaluation reveals that SuperSampler significantly reduces resource usage compared to Sourmash while maintaining similar results. Overall, this work presents a promising approach to making large-scale genomic data more accessible and manageable.

## 2 Preliminaries

This paper presents results on finite strings on the DNA alphabet ∑ = {*A, C, G, T*}, we use *σ* to denote the size of the alphabet. We consider two input multisets of strings longer or equal to *k, S*_*A*_ and *S*_*B*_. These multisets can in practice be read sets from sequencing experiments or genome sequences. We call *k*-mers strings of size *k* over strings of the input sets. *A* = {*x*_0_, *x*_1_, …, *x*_*n*−1_} is the set of distinct *k*-mers from *S*_*A*_ and *B* = {*y*_0_, *y*_1_, …, *y*_*n*−1_} is defined similarly for *S*_*B*_. Our goal is to estimate the similarity between those two sets. Two metrics are mainly used: Jaccard index and the containment index.

### Jaccard Index, containment index and estimators

For two finite, non empty sets of *k*-mers *A* and *B*, the Jaccard index [6] gives a measure of the similarity *A* and *B* by comparing the relative size of the *A* ∩ *B* intersection over the *A* ∪ *B* union,

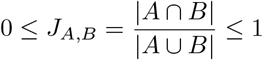

The containment index *C* of the previously defined sets A and B measures the relative size of *A* ∩ *B* intersection over the size of A, i.e., the proportion of distinct *k*-mers of A that are present in B.

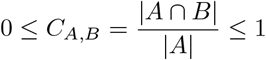

In practice, comparing extremely large sets can be resource-intensive, whereas estimating their similarity can be accomplished much more efficiently. MinHash is a locality-sensitive hashing technique that represents each set with a list of hash values of fixed size. The fraction of shared hash values between two lists provides an estimation of the Jaccard index for the corresponding sets, allowing for efficient similarity measurement.

#### Property 1

(***error in MinHash’s Jaccard estimator****) The error bound E of the MinHash Jaccard estimate* 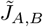 *only relies on the sketch size s [24]*

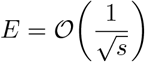

*However, the relative error, E/J*_*A,B*_, *increases significantly for dissimilar documents, as maintaining the error for a decreasing Jaccard index requires a quadratic compensation in the sketch size (refer to additional Figure S1 of [18])*.

Such a property makes fixed-size sketches ill-suited for comparing dissimilar contents, in particular in a resource-frugal context since the sketch size to reach an acceptable precision would be too high to handle.

Happily, scaled sketches that grow linearly with the input size do not present such drawback. Sourmash is the leading tool that implements such scheme and provides sketches composed of *k*-mers represented as 32 bits hashes. To do so, *k*-mers are selected if their hash is below a specific threshold calculated in regard to the desired subsampling rate. Our goal is to improve the space efficiency of such sketch using efficient *k*-mer encoding dubbed SPSS [25] exploiting the fact that *k*-mers share overlaps. The most convenient yet relatively space efficient SPSS are super-*k*-mers and our work relies on such efficient *k*-mer representation using the notion of minimizers.

### Minimizers and super-*k*-mers

#### Definition 2

***(minimizer)*** *Given a total order* 𝒪 *and a k-mer u, the minimizer of u is the smallest m-mer of u according to* 𝒪. *The method to select minimizers is referred to as a minimizer scheme*.

In practice, 𝒪 is defined on integers by hashing *m*-mers using a random hash function *h* and selecting the smallest hash value. We assume that the hash function is chosen such that the hashes are independent and uniformly distributed. From now on, we use *w* = *k* −*m* + 1 to denote the number of *m*-mers inside a *k*-mer.

#### Definition 3

***(super-****k****-mer)*** *A super-k-mer is a maximal substring of a string s (*|*s*| ≥ *k) in which each consecutive k-mers have the same minimizer*.

Super-*k*-mers are an interesting SPSS because a super-*k*-mer containing *x k*-mers is composed of *k* + *x* − 1 bases which incur a cost of 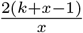 bits per *k*-mer. Therefore longer super-*k*-mers provide lighter *k*-mer representation.

By omitting repeated minimizers inside *k*-mers, the first *k*-mer of the longest possible super-*k*-mers have their minimizers as a suffix. Equally, the last *k*-mers of these super-*k*-mers have their minimizers as a prefix.

#### Definition 4

***(maximal super-****k****-mer)*** *Let s (*|*s*| ≥ *k) be a string and v a super-k-mer of s. Let i*_*m*_ *be the first position of the minimizer on s. v is a maximal super-k-mer iff v starts at position i*_*m*_ + *m* − *k in s and v ends at position i*_*m*_ + *k* − 1 *in s. It follows that v has a length of* 2*k* − *m and contains w* = *k* − *m* + 1 *k-mers*.

Examples of regular and maximal super-*k*-mers are shown in Figure 1. Since maximal super-*k*-mer are the most space-efficient, our approach aims to rely on long super-*k*-mers (ideally maximal) in order to have a compact encoding of the *k*-mer sketch.

**Figure 1.**
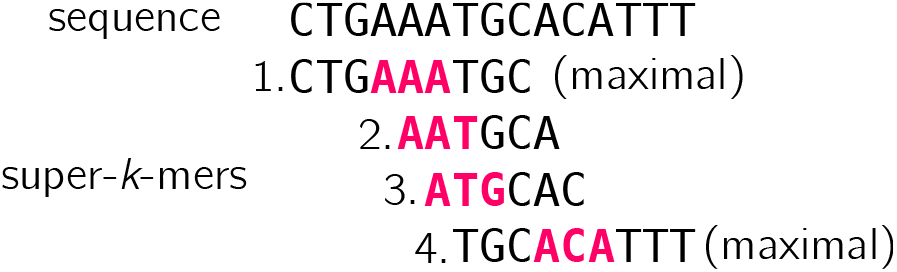
Super-*k*-mers extracted from a sequence for *k* = 6, *m* = 3. Minimizers are shown in purple, here we use the lexicographic order instead of hashing minimizers for the sake of the simplicity. Super-*k*-mers 1 and 4 are maximal (they contain respectively *k*-mers *{CT GAAA, T GAAAT, GAAAT G, AAAT GC}* and *{T GCACA, GCACAT, CACAT T, ACAT TT}*, while 2 and 3 are not (and contain respectively *k*-mers *{AAT GCA}* and *{AT GCAC}*.

#### Property 5

(***coverage of maximal super-****k****-mers****) In [22], authors show that maximal super-k-mers cover approximately half of the k-mers of an input string (Proof of Theorem 3) for a sufficiently large m*.

### Density and universal hitting sets

Since every minimizer corresponds to one super-*k*-mer, the proportion of *m*-mers chosen as minimizers is exactly the inverse of the average length of super-*k*-mers. This proportion, which quantifies the sparsity of a minimizer scheme, is referred to as the *density*.

#### Definition 6

***(density of a minimizer scheme)*** *The density of a minimizer scheme is the expected number of selected minimizers divided by the total number of m-mers. The density factor is equal to the density multiplied by w* + 1.

The density of a minimizer scheme is lower bounded by 1*/w* since each *k*-mer contains *w m*-mers and one of them must be selected as a minimizer. It is known that the expected density of a minimizer scheme based on a random ordering is 2*/*(*w* + 1), and that minimizer schemes cannot have a density below 1.5*/*(*w* + 1) [28]. More generally, *m*-mers selection scheme able to cover every *k*-mer are called *universal hitting sets*.

#### Definition 7

***(universal hitting set or UHS)*** *A set* ∑_*m*_ *is defined as a Universal Hitting Set (UHS) if it includes a minimum of one element from every sequence of w consecutive m-mers*.

Note in particular that the set of all minimizers of **𝒰** ⊆ ∑_*k*_ forms a UHS. Recent publications introduced different methods based on UHS to bring the density below 2*/*(*w* + 1) and closer to the 1.5*/*(*w* + 1) barrier [31, 20]. Thus, a question that naturally arises is: can we cross this barrier by relaxing some constraints on the selection scheme?

## 3 Fractional Hitting Sets

In this section, we introduce the concept of *fractional hitting sets*, which are a generalization of universal hitting sets. These sets are designed to cover a fraction *f* of the *k*-mer space.

### Definition 8

***(fractional hitting set or FHS)*** *A set* ℱ ⊆ ∑_*m*_ *is a Fractional Hitting Set (FHS) if it contains at least one element from a fraction at least f of the sequences of w consecutive m-mers*.

To avoid selection bias in practice, we aim to ensure that *k*-mers are selected randomly, using a random seed as a basis, and that every *k*-mer has an equal chance of being chosen. We introduce a simple way to build such fractional hitting sets by selecting minimizers with a hash smaller than a certain threshold. We call such selected minimizers *small minimizers*. Note that any method selecting a fraction of the minimizers hashes would be suitable here.

### Definition 9

***(small*** *m****-mer)*** *Given a fixed threshold t* ∈ ⟦0, *σ*^*m*^⟧, *we say that a m-mer is small if its hash is below t. We denote by* ***𝒮*** *the set of small m-mers, and* 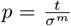 *the probability that a m-mer is small*.

From *p* we can derive the proportion of covered *k*-mers.

### Property 10

*The fraction of k-mers with distinct m-mers containing a small m-mer is*

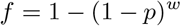

*where w* = *k* − *m* + 1 *and p is the probability that a m-mer is small*.

Proof. Given a *k*-mer with *w* distinct *m*-mers *x*_1_, …, *x*_*w*_,

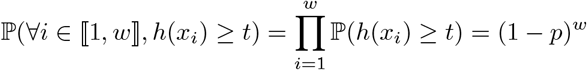

because the hashes of distinct *m*-mers are independent, so

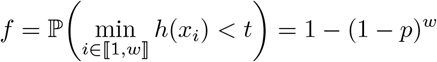

Conversely, if we want a given fraction *f* of the *k*-mers to be covered, the threshold should be chosen as

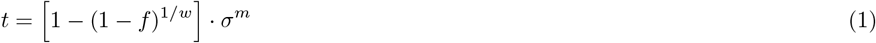

### Density of small minimizers

We showed that selecting *k*-mers with a small minimizer induces an FHS. By considering the usual definition of density (from Definition 6) for this scheme (which cover a fraction *f* of all *k*-mers), we obtain the following bound (proven in supplementary materials):

#### Theorem 11

*Assuming m >* (3 + *ε*) log_*σ*_ *w, the expected density of small minimizers in a random sequence is upper bounded by*

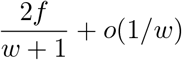

At first glance, the results may be surprising, as the density is smaller than the lower bound of 1*/w* for *f <* 1*/*2 and can approach zero. This is because some *k*-mers may not contain any small minimizers and are therefore not covered, and the proportion of such *k*-mers increases as *f* approaches 0. However, it’s worth noting that this bound does match the 2*/*(*w* + 1) density when *f* = 1 (i.e., when every *k*-mer is covered).

To obtain a more meaningful metric, we can compute the density on the fraction of the sequence that is covered, instead of the entire sequence. With this approach, we obtain the following theorem, which has been proven in the supplementary materials:

#### Theorem 12

***(restricted density)*** *If we restrict to sequences in which every k-mer contains a small minimizer, assuming m >* (3 + *ε*) log_*σ*_ *w, the expected density of small minimizers is upper bounded by*

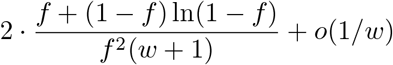

Although less intuitive than the previous one, this result provides valuable insights into the density within the covered portion of the sequence. As shown in Figure 5 (see supplementary materials), the associated density factor ranges from 2 when *f* = 1 (consistent with existing results) to 1 when *f* = 0. Therefore, as *f* approaches 0, we can approach the optimal density.

### Maximal super-*k*-mers

Although measuring the density provides an overview of the average length of super-*k*-mers, it doesn’t indicate how many of them are maximal (i.e., of length 2*k* − *m*). The following result (proven in the supplementary materials) answers this question:

#### Theorem 13

*In a random sequence, the average proportion of maximal super-k-mers (with respect to all super-k-mers) built from small minimizers is given by*

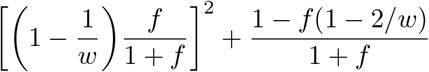

Note that this result generalizes theorem 4 from [22], which corresponds to *f* = 1. As shown in figure 12b (see supplementary materials), the proportion increases towards 100% as *f* approaches 0.

### Improving the density of fractional hitting sets using UHS

This effect is more pronounced for smaller values of *f*. This observation raises a natural question: what is the lowest achievable density for a given *f*? Since universal hitting sets (UHS) with a density lower than 2 have been proposed for *f* = 1, it is possible that they may also improve the density for smaller *f* values by considering only the *m*-mers selected by the UHS as potential minimizers.

#### Theorem 14

*Given a UHS* **𝒰** *with density d*_**𝒰**_, *assuming m >* (3 + *ε*) log_*σ*_ *w, the expected density of small minimizers selected from* **𝒰** *(that is*, **𝒮** ∩ **𝒰***) in a random sequence is upper bounded by*

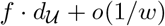

The proof is given in supplementary materials. Note that this result generalizes theorem 11 since the UHS of minimizers selected using a random ordering has a density of 2*/*(*w* + 1) [28].

## 4 SuperSampler

### 4.1 Related work

#### Sketching

Sketching strategies involve building a summary, or “sketch”, of an input dataset that is smaller than the original set. The purpose of this sketch is to approximate the similarity between two documents based on the similarity between their respective sketches. In our context, sketches are lists or sets of fingerprints that represent uniformly selected *k*-mers from the set.

#### Fixed-size sketches

MinHash is a seminal method for fixed-size sketch-based set comparison. In our context, a MinHash sketch is a list of *k*-mer hashes that are minimal in some sense. As these *k*-mers are selected uniformly, the proportion of shared hashes between two sketches approximates the Jaccard index of their original documents. Tools such as MASH [18] employ this principle to efficiently compare large collections. However, two main limitations arise from this strategy. First, hash collisions need to be addressed, as they introduce more resemblance between two compared sets than what actually exists. Therefore, the fingerprint size must be chosen appropriately to control such collisions. Recent approaches like [30], Dashing [2], Dashing2 [3] and NIQKI [1] used enhanced fingerprint scheme related to Hyperloglog [7] or to HyperMinHash [29] to get the best possible precision memory trade-off. Second, as stated in Property 1, this approach is best suited for comparing sets of similar size and content. When comparing sets with significant size or content differences, the accuracy of the estimate decreases substantially.

#### Scaled sketches

As the problem of inaccuracy of fixed-size sketches for dissimilar datasets was pointed out but only partially solved using MinHash schemes [17, 13], novel approaches were proposed. Scaled sketches methods build their sketches with adapted sizes in regard to the input. The main contribution in this direction for compositional analysis is sourmash [23], and our paper also falls in this category. sourmash incorporates a scaled factor indicating the fraction of the *k*-mer’s input that should be included in the sketch as fingerprints. To do so they add a fingerprint in the sketch if their integer value is a multiple of *S* (ScaledMinhash) or lower than a fixed threshold (FracMinHash). The latter technique is recently described and theoretically studied in [10, 8]. In contrast to previously presented fixed-size sketching approaches, those approaches have the advantage of providing a more accurate comparison of uneven sized sets. This flexibility comes at the cost of possibly large sketches. Schemes based on minimizers or other seeds are also used for read mapping [11, 27] but are out of the scope of this paper.

### 4.2 SuperSampler’s sketch construction

#### Definition 15

(***SuperSampler’s sketch****) Given a sequence S, each super-k-mer whose minimizer’s hash is lower or equal to a threshold T is selected, and all its surrounding k-mers are kept in the sketch as a super-k-mer. A supersampler sketch can therefore be represented as a super-k-mer set*.

In Sourmash, *k*-mers are represented as integer fingerprints through hashing (using a default of 32-bit). This approach reduces space requirements compared to employing 2*k* bits per *k*-mer, but it also introduces the potential for false positives due to hash collisions. However, the false positive rate is exponentially low, depending on the fingerprint size, which allows for efficient control.

In contrast, SuperSampler explicitly stores *k*-mers as super-*k*-mers, offering two significant advantages over the conventional method: First, the selected *k*-mers are represented exactly, eliminating false matches and enabling the output of shared *k*-mers when they are of interest to users. Second, this technique enables a more space-efficient representation of *k*-mers, typically requiring less than the 32-bit per *k*-mer space cost of Sourmash. Figure 2 illustrates an example of sketch construction in SuperSampler. In the following sections, we propose a model to evaluate the space efficiency of a SuperSampler sketch.

**Figure 2.**
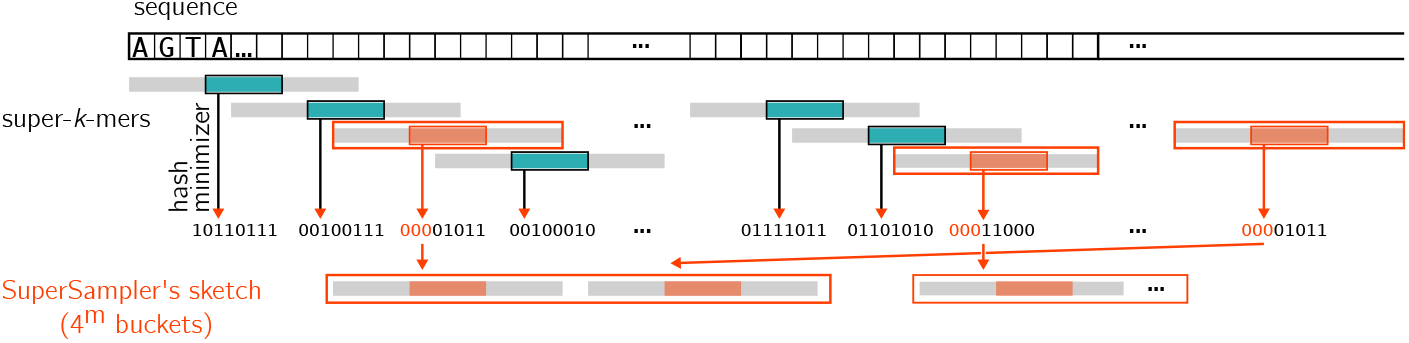
SuperSampler’s sketching strategy. In order to build sketches, SuperSampler computes super-*k*-mers over the input sequence. Fingerprints are associated with each super-*k*-mer by hashing their minimizers to an integer, hence an integer per super-*k*-mer. Super-*k*-mers associated to sufficiently low integers are kept in the sketch. Super-*k*-mers are put into buckets according to their minimizer.

In the regular case with hashed minimizer with a density factor of 2 we can expect (*k*− *m* + 1)*/*2 *k*-mers per super-*k*-mers [22]. This results in a mean super-*k*-mer length of (3*k* −*m* −1)*/*2 bases. We can give a lower bound of the cost in bit per *k*-mer to encode such super-*k*-mers:

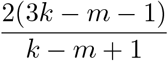

However we need to encode the length of each super-*k*-mer to avoid considering artefactual *k*-mers created by two successive super-*k*-mers so we can add log_2_(*k* − *m* + 1) bits per super-*k*-mer (encoding the number of *k*-mers). This leads to a bits per *k*-mer ratio of

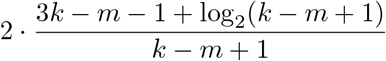

In practice we use the formula (1) to select a *k*-mer fraction chosen by the user. As a side effect, since we used an FHS, selected super-*k*-mers are longer than those selected by regular hashed minimizer scheme as our hitting set provides a lower density. Importantly for low selected fraction a very large proportion of super-*k*-mers are maximal. This property is of prime interest because maximal super-*k*-mers can be efficiently encoded for two reasons. First they are all of the same size so we do not need to encode their respective length or any kind of separator. Second, they represent 2*k* − *m* bases encoding for *k* − *m* + 1 *k*-mers, they provide a lower bits per *k*-mers ratio

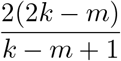

### 4.3 Partitioned sketches

Minimizers naturally partition the super-*k*-mer space into **𝒪**(4_*m*_) buckets. Since only a subset of the minimizer space is selected, a smaller number of buckets are actually considered. SuperSampler relies on the fact that super-*k*-mers are centered around a shared minimizer to build a partitioned sketch. In practice, we examine all non-empty selected buckets, each storing a minimizer with their corresponding super-*k*-mers independently. This strategy offers several crucial advantages. First, when encoding maximal super-*k*-mers, we know the position of the minimizer within each super-*k*-mer. By storing the minimizer sequence once in the bucket, we can omit it in all maximal super-*k*-mers. This results in an even lower bit per *k*-mer ratio:

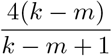

Figure 3 shows the space cost of the different encoding: super-*k*-mer, maximal super-*k*mer and partitioned super-*k*-mer according the the minimizer size along with the actual performances of Supersampler. While this mechanism enhances space efficiency, it also significantly improves actual memory usage. When comparing document sketches, matching *k*-mers between documents are necessarily found in the same partition, so only a given partition is needed in memory at a time. For sufficiently large *m*, such partitions should be orders of magnitude smaller than the total amount of selected *k*-mers, as the expected partition size decreases exponentially with *m*. This partitioning technique also allows for substantial speed-ups, notably in sketch comparison time, which are discussed later on.

**Figure 3.**
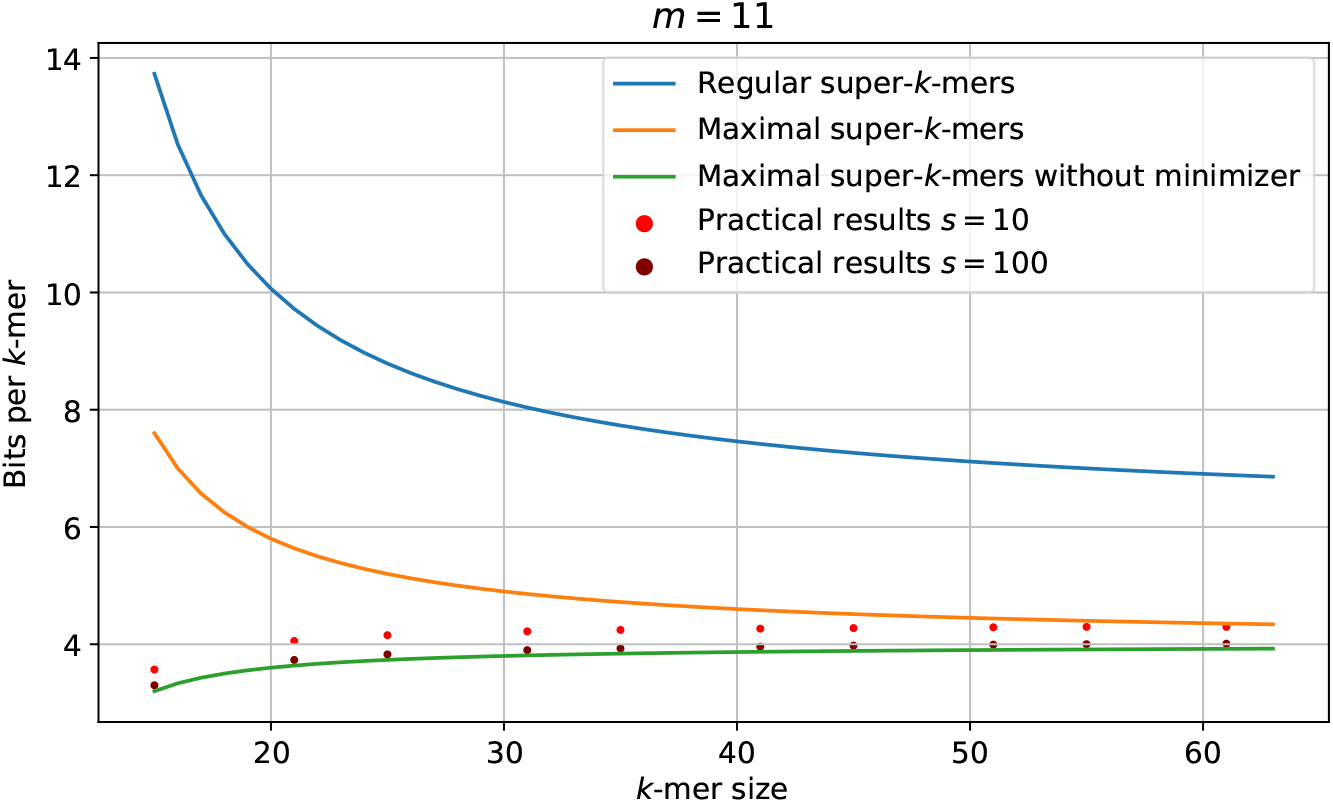
Theoretical space cost of different encodings in bits per *k*-mer according the *k*-mer size size along with practical space usage of super-sampler sketches on random sequences.

### 4.4 Sketches comparisons

#### Sketches comparisons

Unlike sourmash that treats sketches in their entirety, SuperSampler focuses on small related partitions. This allows for two distinct computational improvements. First, a partition that is specific to a file can be skipped as we know that no *k*-mer present in such partition will be found in another file so no matching *k*-mer can be found. Second, the size of the buckets stored in memory being small, we expect few cache-misses when comparing partitions unlike sourmash for which a cache-miss for each queried fingerprint can be expected. In other words, as illustrated in Figure 4, SuperSampler concentrates exclusively on small, relevant partitions and processes each of them only once. The efficiency benefits of this approach are magnified when comparing one or multiple documents against a large collection, as SuperSampler processes only a specific partition of the relevant documents at a given time. This targeted processing reduces the computational load and enhances the overall performance of the comparison.

**Figure 4.**
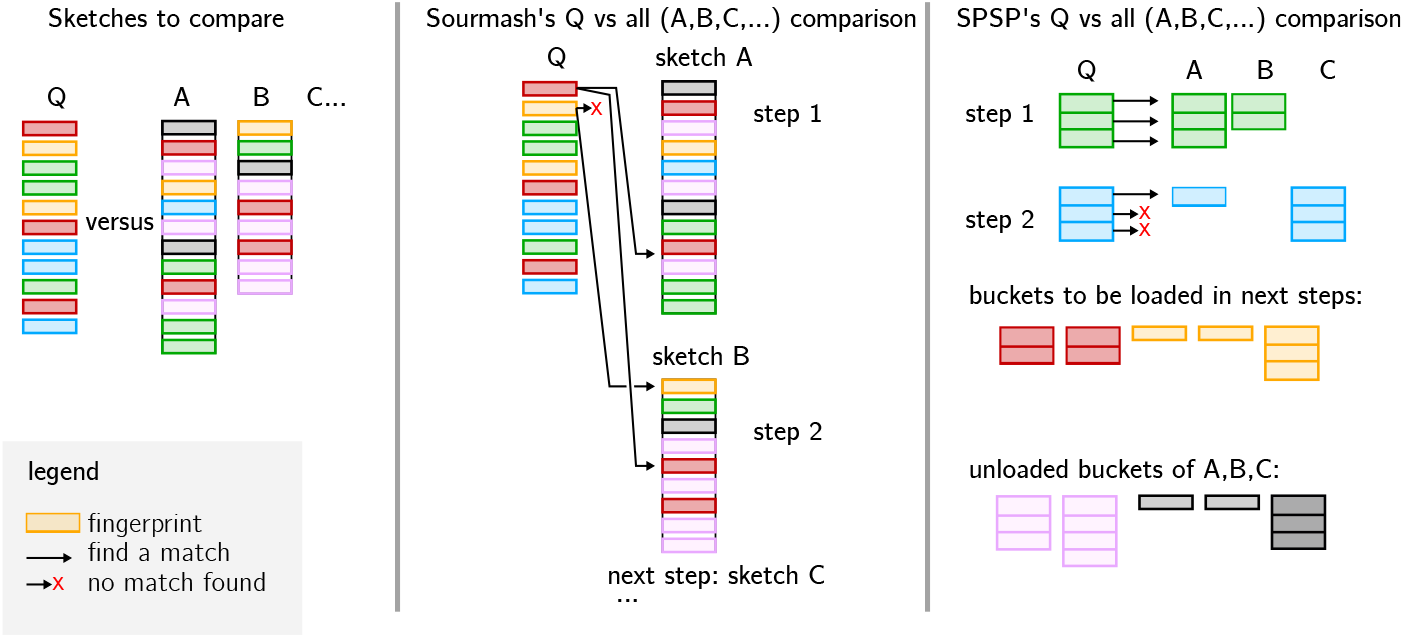
How SuperSampler and Sourmash perform their respective sketch comparison. In this example, we discuss the comparison of one document against a collection, although other use cases can be inferred. SuperSampler is capable of skipping certain buckets that are not relevant to the query. By focusing on smaller sub-parts of the collection one at a time, SuperSampler effectively improves practical performance and reduces memory usage.

## 5 Results

All experiments were performed on a single cluster node running with Intel(R) Xeon(R) Gold 6130 CPU @ 2.10GHz and 2 64GiB DIMM DDR4 Synchronous 2666 MHz ram.

In the first section, we evaluate supersampler sketches space usage. In the second section, we evaluate the precision and performance of SuperSampler in comparison to Sourmash, the current state-of-the-art solution. Finally, in the last section, we demonstrate SuperSampler’s scalability when indexing extensive collections. Every file used to perform these experiments can be found GitHub : github.com/TimRouze/Expe_SPSP.

### 5.1 Space efficiency of supersampler

As previously discussed, the lower bound of memory cost for storing *k*-mers as super-*k*-mers is given by

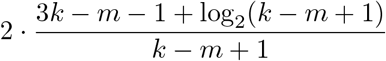

bits per *k*-mer, assuming we store the size of each super-*k*-mer in the index. For *m* = 15, this bound equates to 9.2 bits/*k*-mer with *k* = 31 and 7.1 bits/*k*-mer with *k* = 63. However, in practice, SuperSampler exhibits lower space usage. Figure 13 in the supplementary materials reveals that approximately 6.5 bits/*k*-mer and 5 bits/*k*-mer are achieved for *k* = 31 and 63, respectively.

These results can be attributed to the low density permitted by SuperSampler’s minimizer selection scheme. As illustrated in Figure 5, the density factor quickly diminishes as the subsampling rate increases, respectivey as the fraction f diminishes. When the subsampling rate is 2, the density factor falls below 1.5, the lower bound of the minimizer scheme, and continues to decline toward 1. In general, subsampling tools seldom apply rates below 100, with Sourmash defaulting to a rate of 1000 i.e. *f* = 1*/*1000. Consequently, SuperSampler consistently remains close to the lower bound for the density factor, since the density factor for a subsampling rate of 100 is already below 1.04. This facilitates the indexing of longer super-*k*-mers, which are stored more efficiently as their length increases.

**Figure 5.**
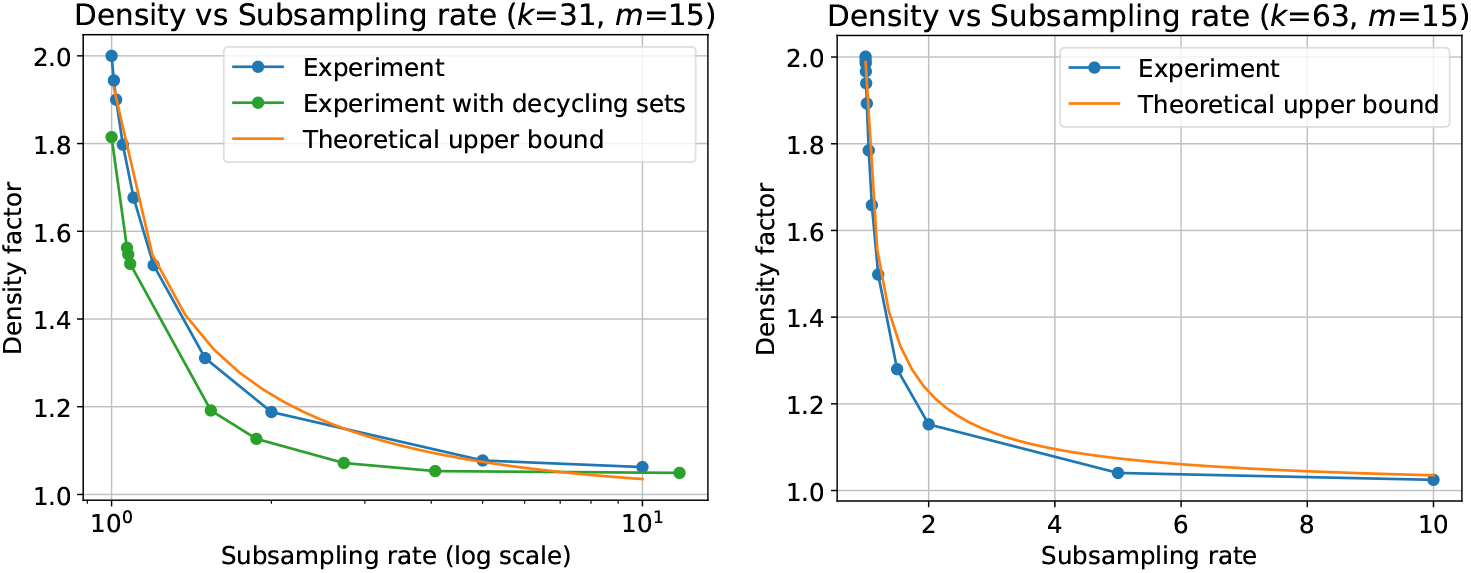
Measured density factor compared to the model.

To further reduce memory costs, SuperSampler offers an option to use its selection scheme in conjunction with existing UHS-based minimizer schemes, specifically the modified double decycling sets introduced in [20]. As depicted in Figure 6, this approach marginally improves the bit/*k*-mer cost, particularly for smaller values of *k*.

**Figure 6.**
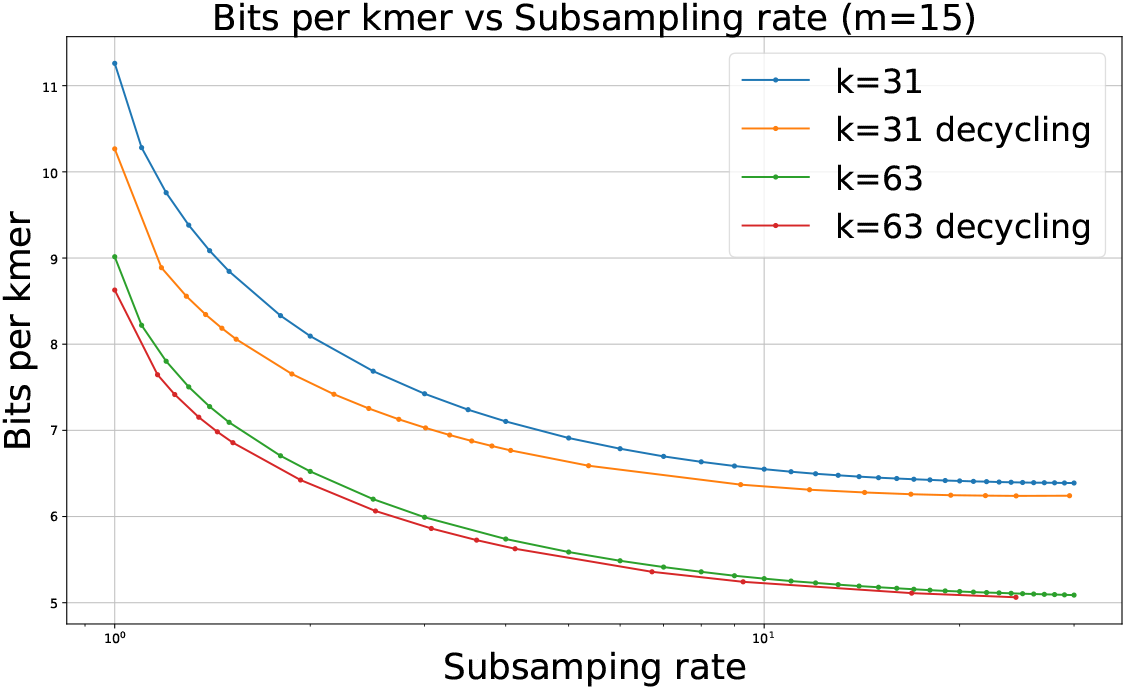
Space cost in bits per *k*-mer according to the subsampling rate with and without using the improved UHS.

A high proportion of maximal super-*k*-mers in a sketch is advantageous for SuperSampler, as it lowers the bit/*k*-mer cost. Figure 7 demonstrates that the percentage of maximal super-*k*-mers increases rapidly with the subsampling rate, reaching 90%, 99%, and 99.9% of indexed super-*k*-mers when the subsampling rate is around 10, 100, and 1000, respectively. This feature is particularly significant because it enables a rapid and considerable reduction in the bit/*k*-mer cost by efficiently encoding maximal super-*k*-mers. Therefore, with the subsampling rates commonly used in practice, which involve a very high proportion of maximal super-*k*-mers, the actual bound is determined by the following formula

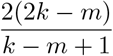

**Figure 7.**
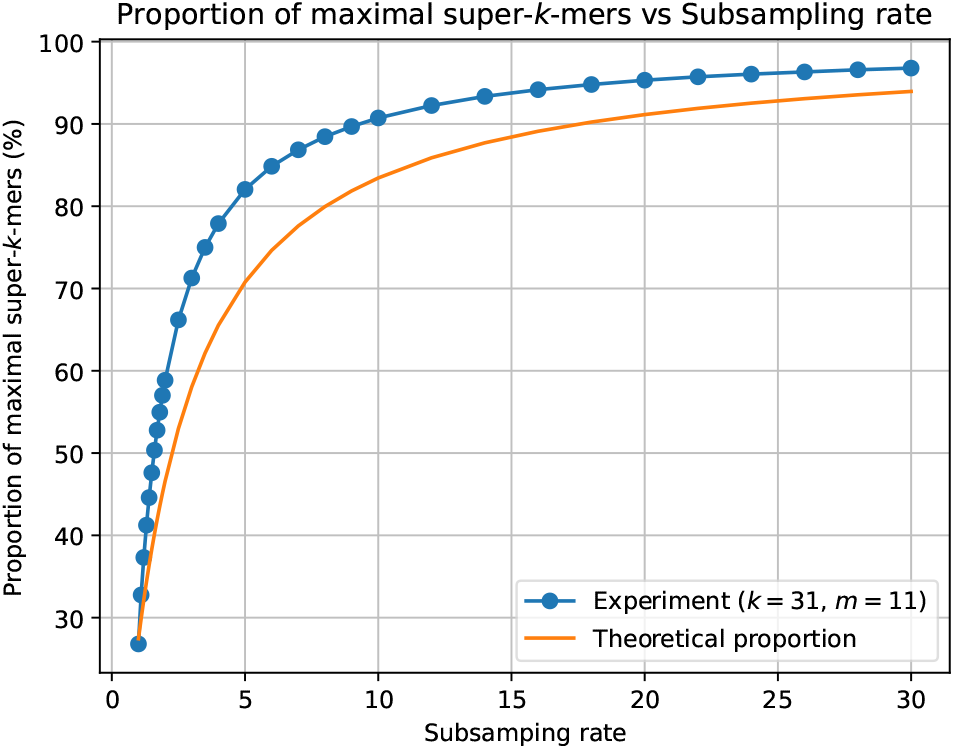
Fraction of maximal super-*k*-mers according to the subsampling rate.

### 5.2 Performance Comparison

In our qualitative experiment, we compared the performance of SuperSampler with Sourmash, which implements FracMinHash. We evaluated both tools on two distinct datasets: 1024 Salmonella genomes from GenBank and 1024 bacterial genomes from RefSeq. These collections were chosen due to their differing containment indexes; Salmonella genomes are highly similar to each other, while the bacterial genomes in RefSeq exhibit much greater dissimilarity (Jaccard index close to 0).

We carried out an all-versus-all comparison of these collections using both tools and monitored RAM and disk usage, as well as computation time during the sketch comparisons. To assess the precision of the approximated scores, we calculated the exact Jaccard and containment index values using Simka [4], which performs efficient *k*-mer counting operations on large collections. With these scores as a reference, the precision of the approximation can be evaluated.

RAM and computation time were measured using the benchmark flag from Snakemake, with one run per command. Disk usage was determined by comparing the sketch sizes of Sourmash (using the *zip* option for *sourmash sketch*) and SuperSampler, with the latter’s sketch sizes examined through a Python script. SuperSampler sketches were stored in a tar archive and compressed using *gzip -9*.

#### 5.2.1 RAM, time, and sketch size

Figures 14 and 8 demonstrate that, for *k* = 31, SuperSampler generally consumes 5 times less RAM and requires generally 16 times less space than Sourmash. Additionally, SuperSampler performs computations 50 times faster than Sourmash when comparing highly dissimilar genomes. However, when genomes are very similar, such as with Salmonella, comparison times are comparable since SuperSampler’s time optimization does not apply on very similar documents. We also note that minimizer size has little to no impact on these metrics.

**Figure 8.**
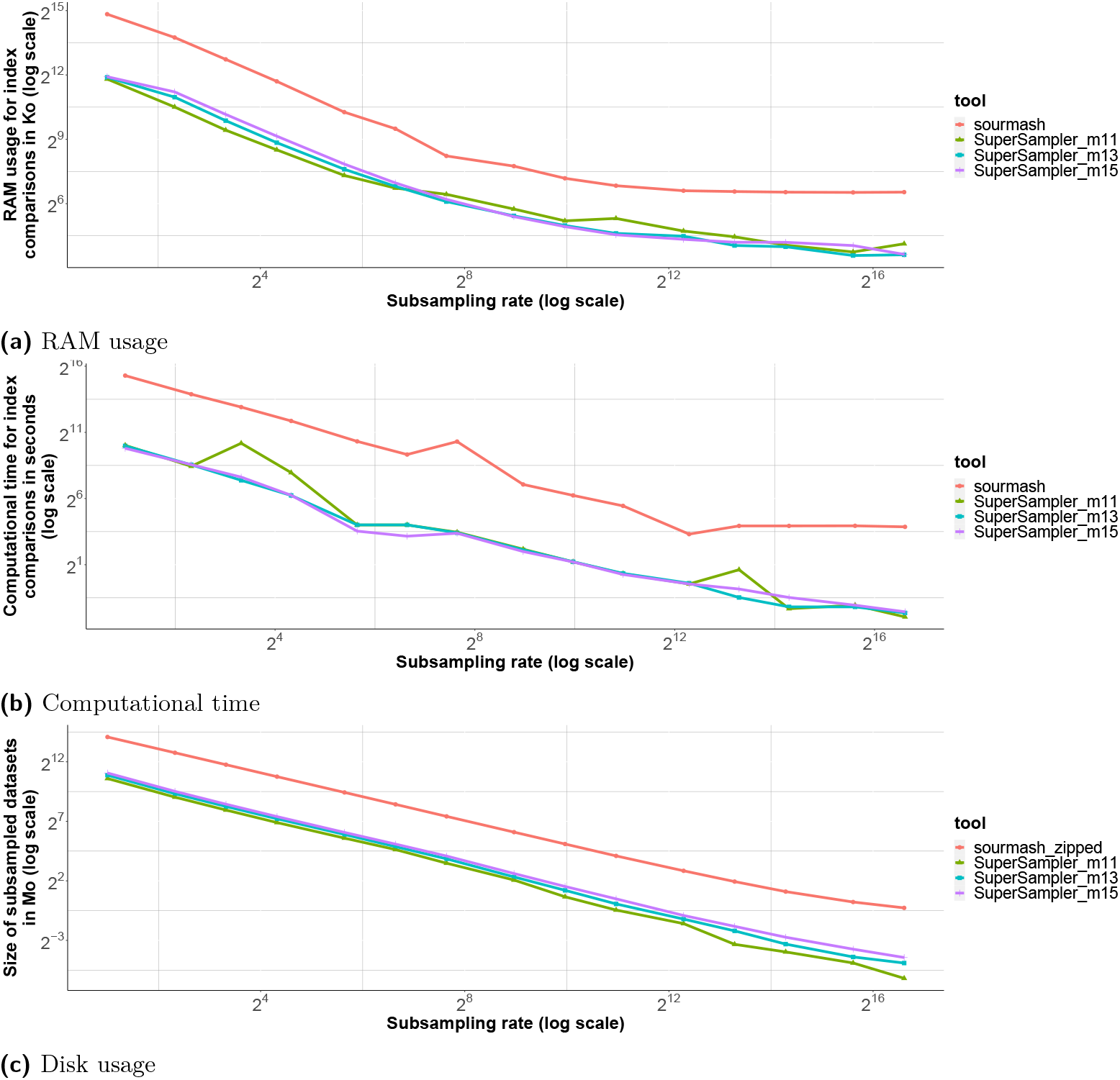
Resource consumption results for 1024 bacterial genomes from RefSeq. For these results, k = 31. For results on Salmonella genomes and k = 63, see figures 14, 16 and 15 on the appendix.

Figures 15 and 16 reveal that the improvement in sketch disk size is even more significant with larger values of *k*. SuperSampler uses in general 50 times less disk space than Sourmash with *k* = 63, while maintaining similar differences in RAM and computation time.

#### 5.2.2 Error

As demonstrated in Figure 9, SuperSampler’s performance is similar to Sourmash in terms of error, although it exhibits slightly lower accuracy. This reduced accuracy results from a clustering effect caused by SuperSampler selecting overlapping *k*-mers around small minimizers. This effect is counterbalanced by SuperSampler’s ability to index and compare more *k*-mers using the same amount of memory and generally less computation time. Figure 9 shows strong variability in error for both Sourmash and SuperSampler, it is to be noted that the error is constantly small and that small variability on already small number leads to such fuzzy figures.

**Figure 9.**
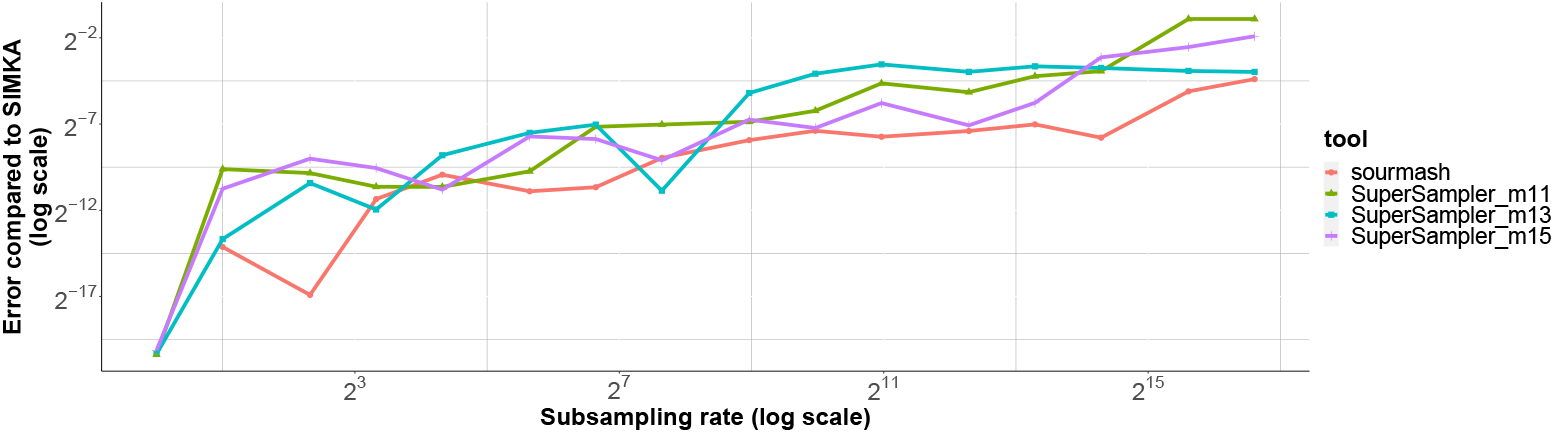
Error against Simka on Containment index approximation for Sourmash (red line) and SuperSampler with different minimizer sizes. This plot is for 1024 Salmonellas genoms with *k* = 63. Other results for RefSeq and Salmonellas are available at figure 17. Jaccard index error is available at figure 18 on the appendix.

Additionally, SuperSampler stores *k*-mers in plain text without any loss of information, which means that *k*-mers of interest can actually be retrieved. While Sourmash could rely on invertible hash functions and larger hashes to match this ability, doing so would effectively increase their space usage.

#### 5.2.3 Massive collection indexing

As a scalability experiment, SuperSampler and Sourmash were monitored on their performances while analysing growing collection of RefSeq bacterial genomes. Result are displayed in Figure 10. We can see that our tool is able to handle all versus all comparison very large scale collection comparison effectively. For the biggest amount of files, SuperSampler took 25 CPU hours. We observe that the gap between Sourmash and Supersampler is diminishing for larger collection as the output matrix itself become large and generate cache-miss for every update. The sketch creation step is essentially cheap as both tools only took a couple CPU hours the actual bottleneck being IO.

**Figure 10.**
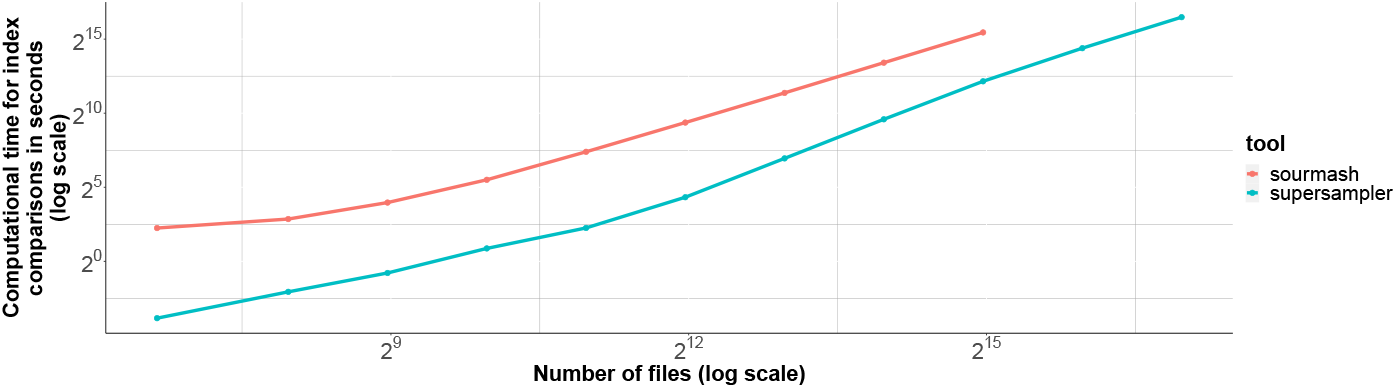
Computational time for comparisons on different amounts of bacterial genomes from RefSeq. From 100 to 128,000 genomes with *k* = 63, *s* = 1000 and *m* = 15 Sourmash was run up to 32,000 genomes as it was taking too much time for the 2 last experiments.

**Figure 11.**
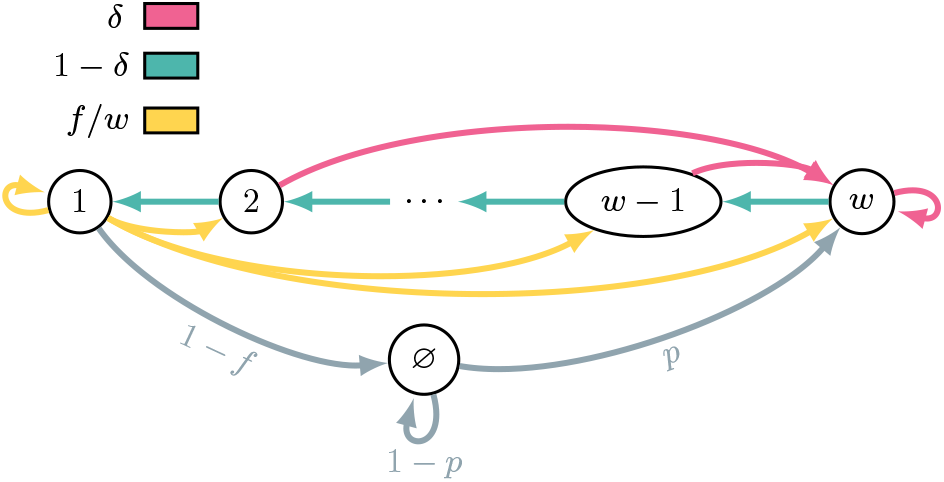
The chain is in state *i* ∈ *⟦*1, *w⟧* if the small minimizer starts at position *i* in the *k*-mer, and ∅ if there is no small minimizer. Different edge colors represent different probabilities.

**Figure 12.**
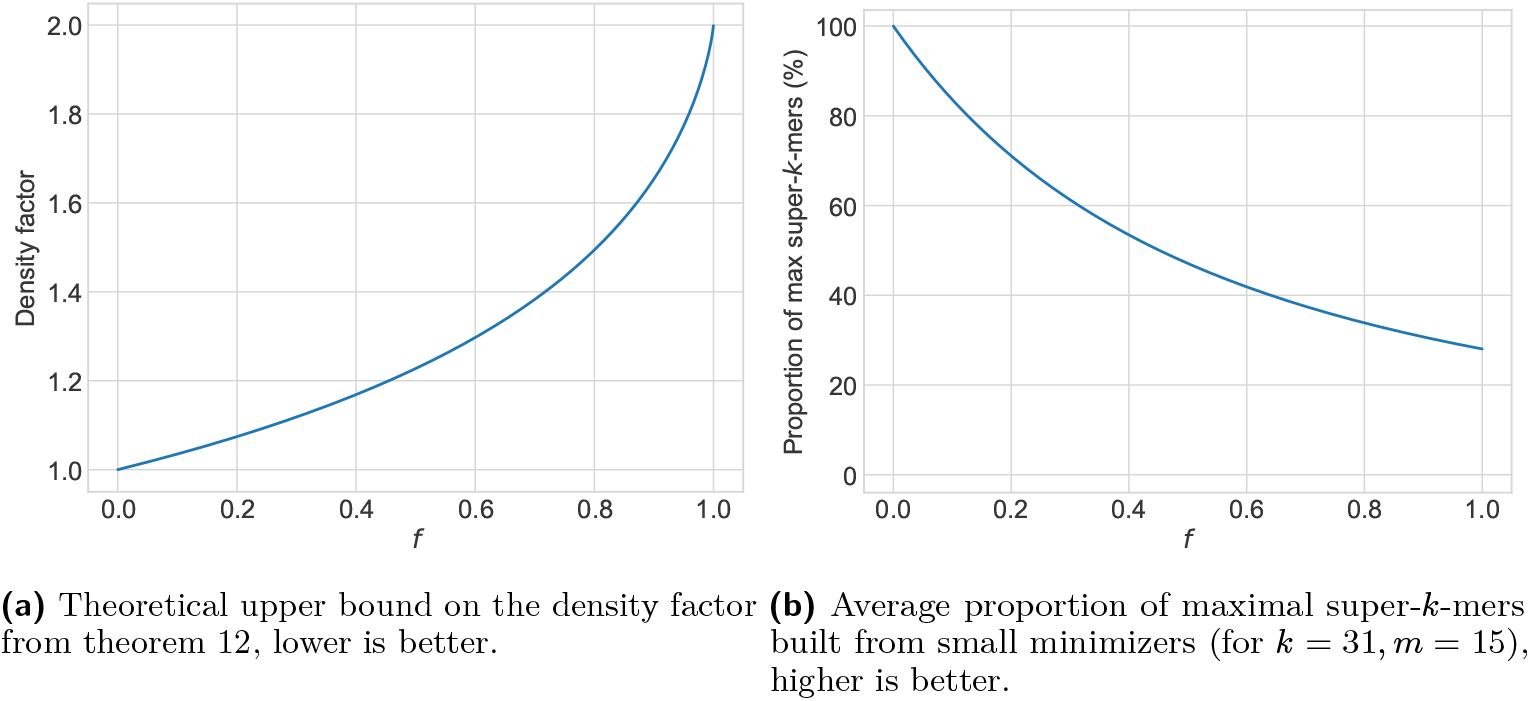
Theoretical bounds on the density factor and the proportion of maximal super-*k*-mers depending on the fraction *f* of covered *k*-mers.

**Figure 13.**
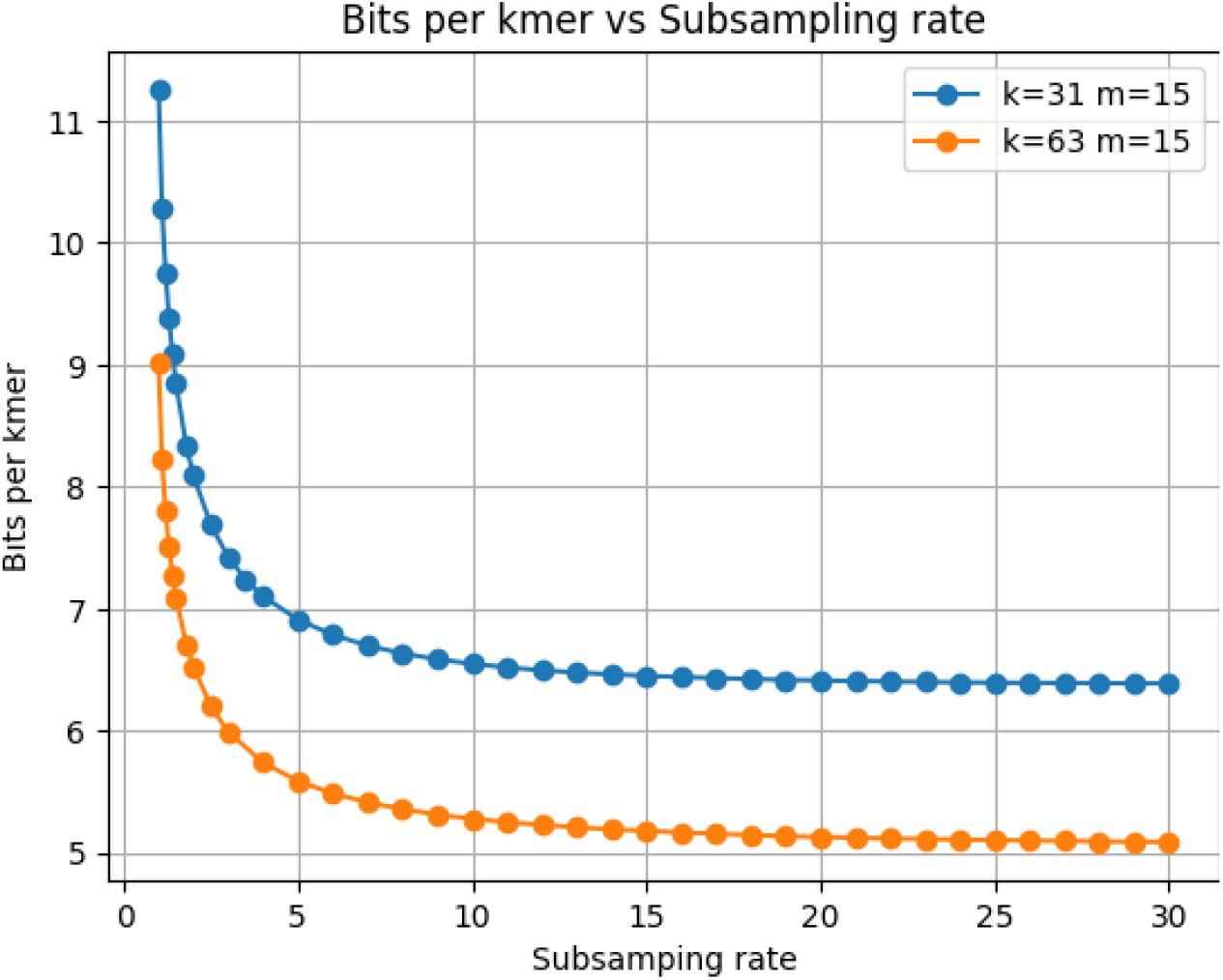
Space cost in bits per *k*-mer according to the subsampling rate.

**Figure 14.**
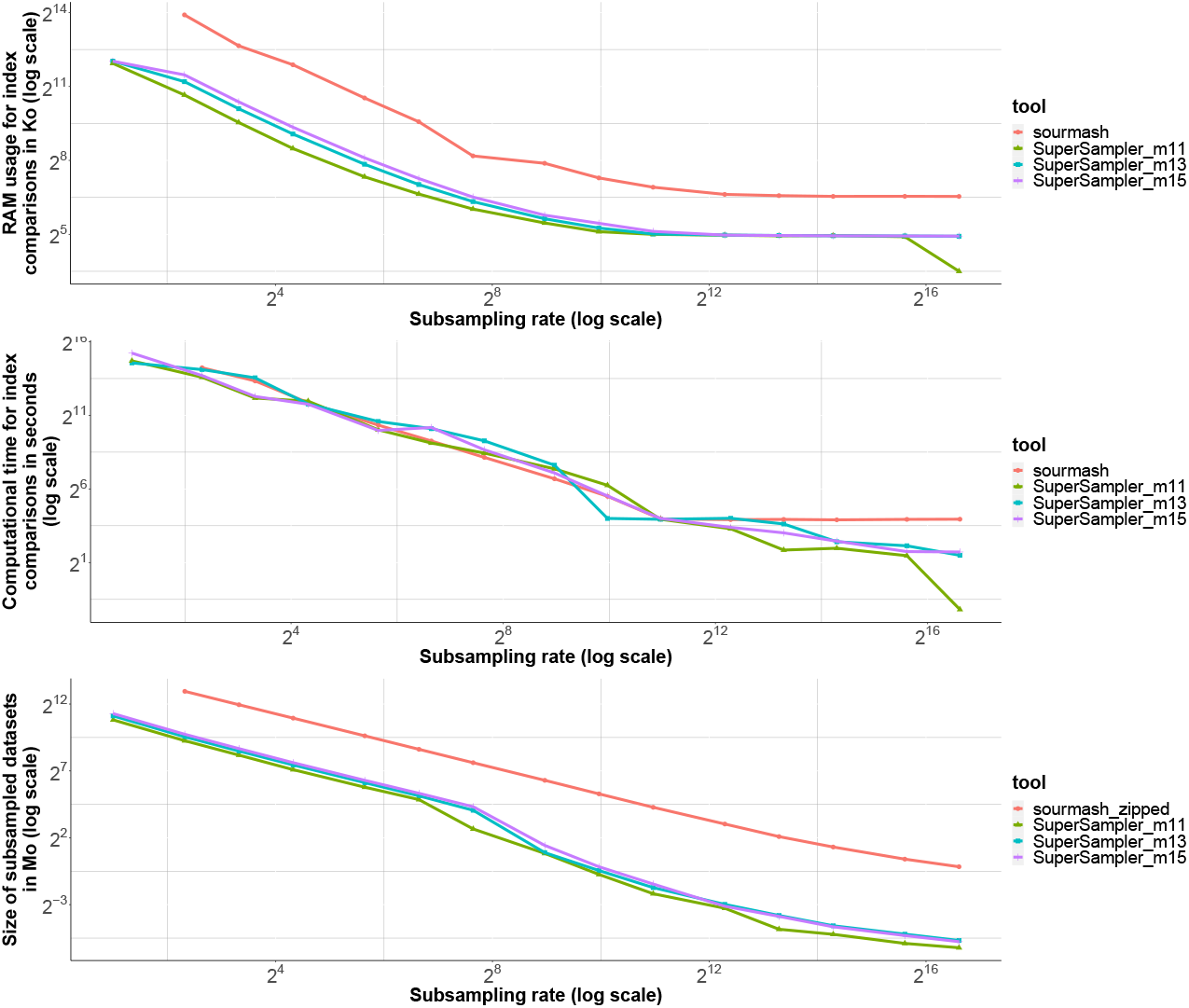
Salmonellas 1K k=31

**Figure 15.**
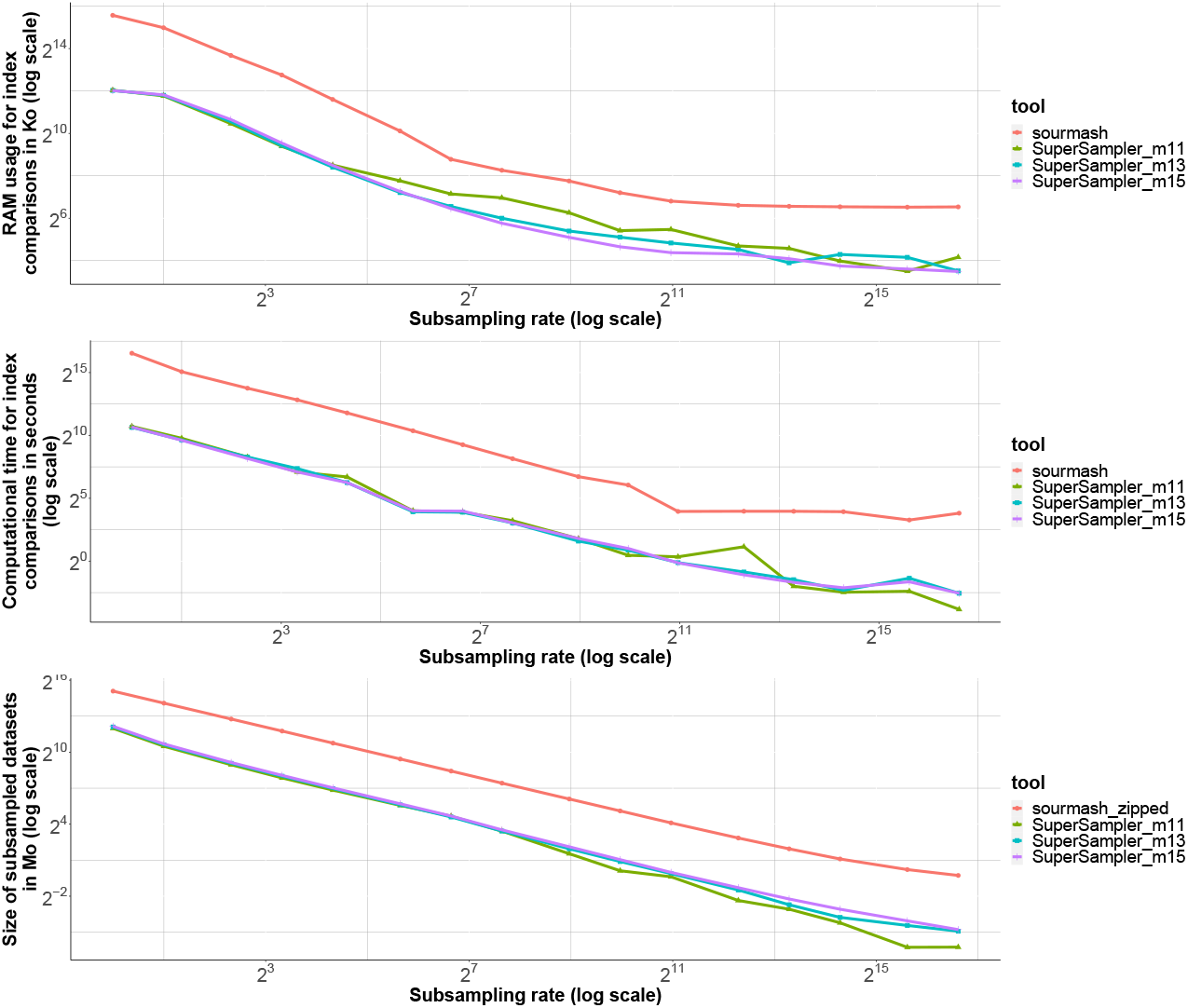
Refseq 1K k=63

**Figure 16.**
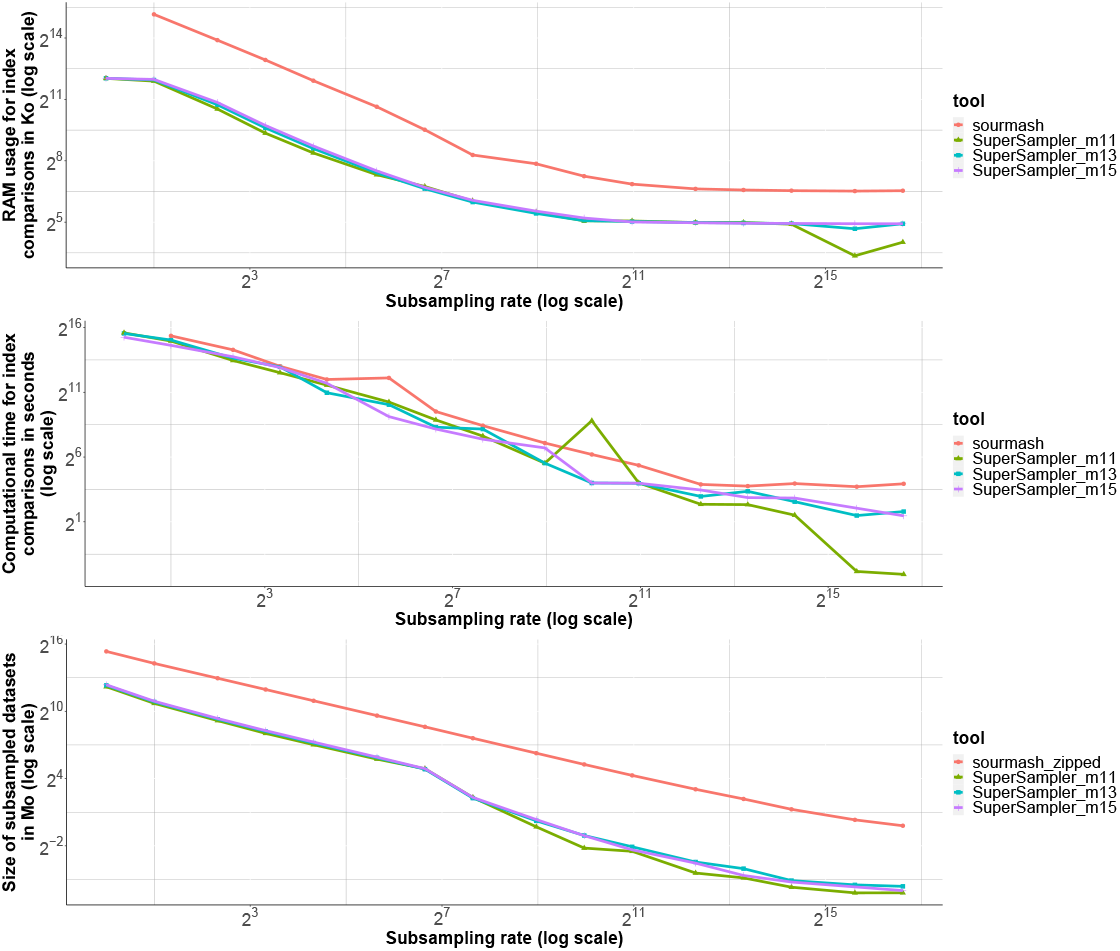
Salmonellas 1K k=63

**Figure 17.**
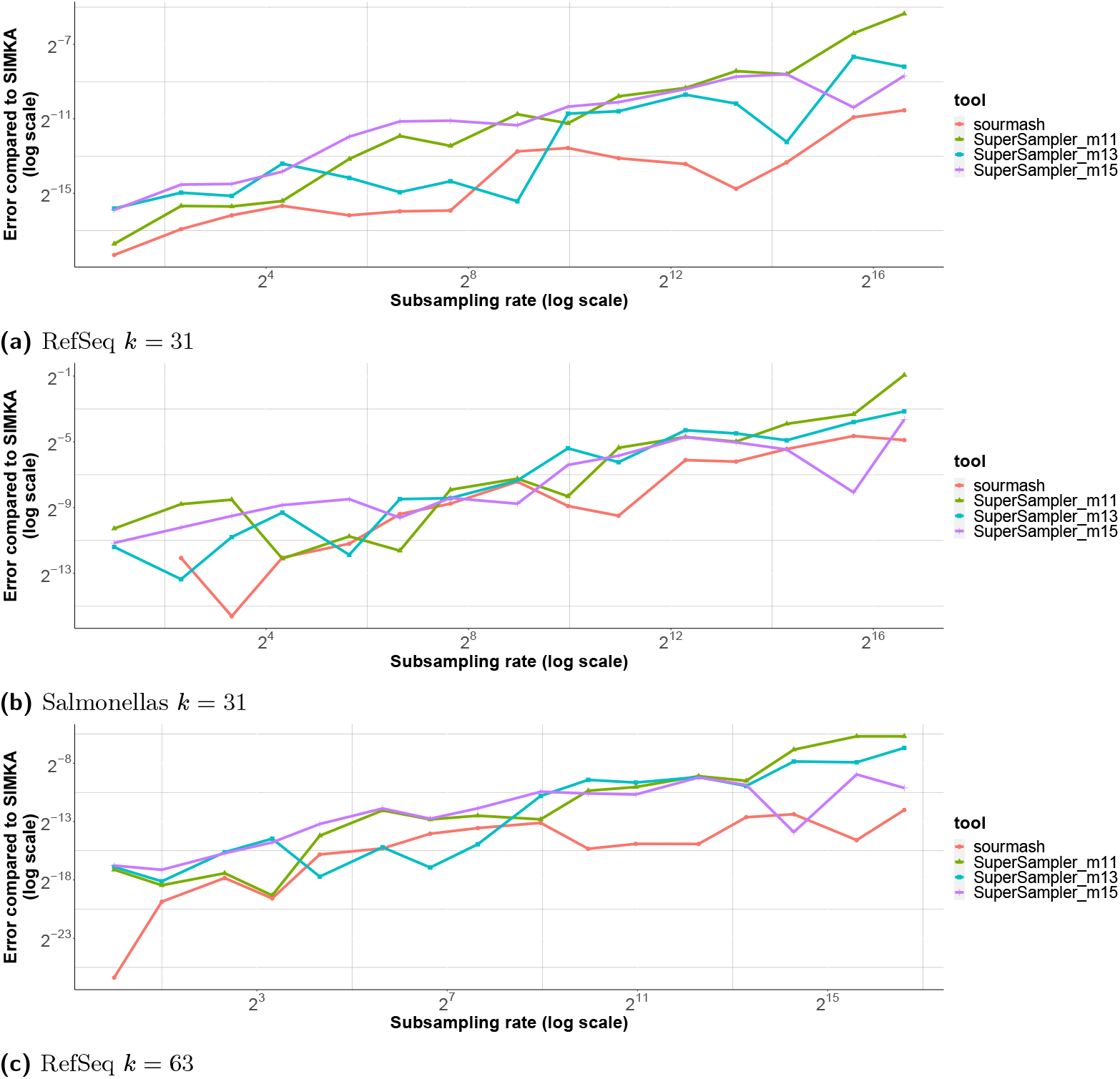
Error for containment index approximation for Sourmash (red line) and SuperSampler on different values for minimizer sizes.

**Figure 18.**
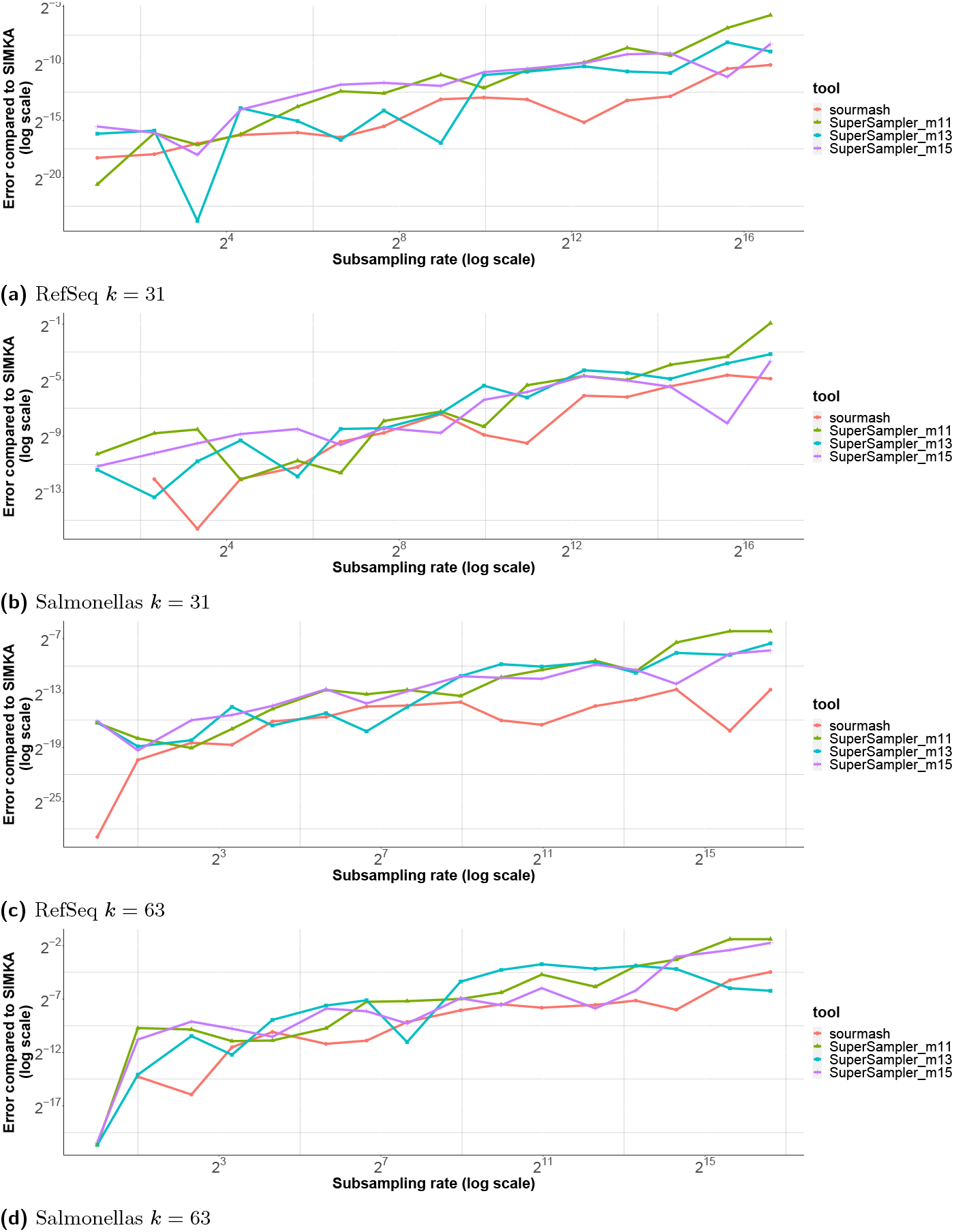
Error for Jaccard index approximation for Sourmash (red line) and SuperSampler on different values for minimizer sizes.

## 6 Conclusion

In this paper, we present both theoretical and practical results of an innovative subsampling scheme based on super-*k*-mers. We introduce the fractional hitting sets framework and propose a straightforward sketching method to highlight its benefits. This approach offers improved density compared to other schemes and tends to select *k*-mers that contribute to better space usage. Capitalizing on this scheme, we propose SuperSampler, an open-source sketching method for metagenomic assessment.

Through comprehensive experimental evaluation, we demonstrate that SuperSampler enables efficient and lightweight analysis of extensive genomic data sets with fewer resource requirements compared to the state-of-the-art tool Sourmash. More generally our results confirm the validity of our methodology from both theoretical and experimental standpoints. We recognize several potential enhancements for our study. First, concerning SuperSampler’s implementation, we aim to refine the tool for increased user-friendliness and adaptability for routine analysis while augmenting its capabilities, such as abundance tracking. Implementation of such improvements will lead to more thorough experiments with existing sampling methods as well as new comparisons with Sourmash using the same amount of disk memory in order to better show SuperSampler’s capacity with regard to both fixed-size and scalable sketches.

Second, we plan to investigate alternative methods for sketch comparison, like sorted fingerprints, which could potentially reduce the complexity of the comparison process. From a theoretical perspective, delving deeper into the properties of Fractional Hitting Sets and gaining a better understanding of density and restricted density bounds for various values of *f* may lead to even more efficient and robust sketching techniques.

## A Useful lemmas

### Lemma 16

*Assuming p is non-increasing with respect to w*, 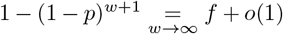

Proof. (1 − *p*)_*w*+1_ = (1 − *p*)_*w*_ − *p*(1 − *p*)_*w*_ if 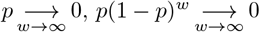 otherwise *p* ≥ *c* for some *c >* 0 since it is non-increasing, so 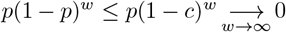

Therefore, (1 − *p*)_*w*+1_ = (1 − *p*)_*w*_ + *o*(1)

We use 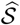 to denote the set of *k*-mers containing a small *m*-mer.

### Lemma 17

*Given two consecutive k-mers W*_0_ *and W*_1_, 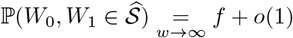

Proof.

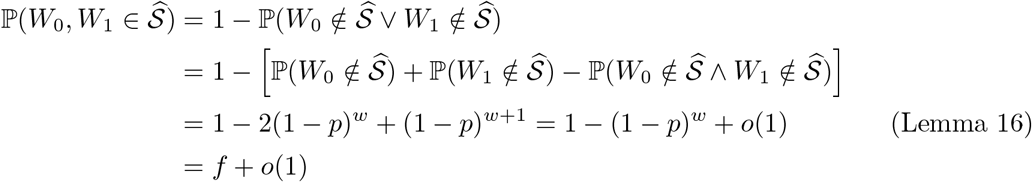

### Lemma 18

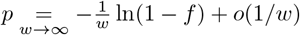

Proof. Because of property 10, we have *p* = 1 − (1 − *f*)_1*/w*_ and

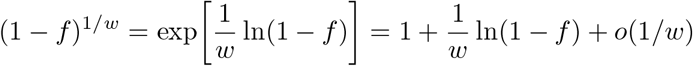

## B Proof of theorem 11

In order to upper bound the density, we follow the same approach as the one presented in [31] (for the proof of theorem 7). As stated in [31], the density is equivalent to the probability that a context *c* (that is, the string formed by two consecutive *k*-mers) is *charged*, i.e. the two *k*-mers of *c* have different minimizers.

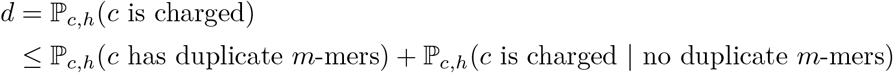

### Lemma 19 (lemma 9 from [31])

*Assuming m >* (3 + *ε*) log_*σ*_ *w*,

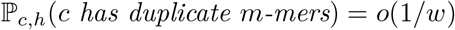

If *c* has no duplicate *m*-mers, the small *m*-mers are all distinct and each of them has the same probability to be minimal since *h* is random. Therefore,

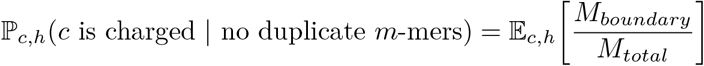

where *M*_*boundary*_ denotes the number of boundary *m*-mers that are small and *M*_*total*_ denotes the total number of small *m*-mers in *c*.

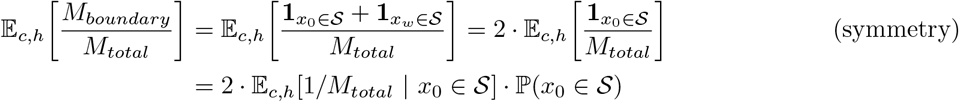

Assuming *x*_0_ is small, we have *M*_*total*_ = 1 + *X* with *X ∼ B*(*w, p*), since each other *m*-mer of *c* has a probability *p* to be small.

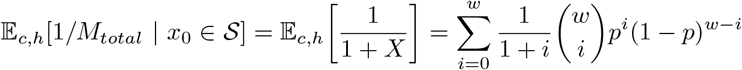

### Lemma 20

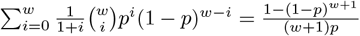

Proof.

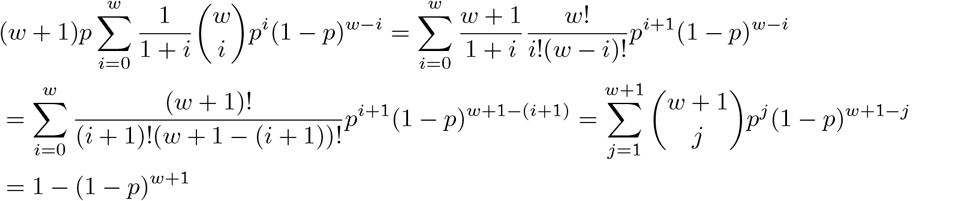

Finally, since 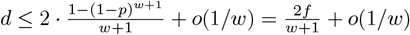 (Lemma 16)

## C Proof of theorem 12

In this section, **we assume that every** *k***-mer we work with contains a small** *m***-mer**.

Just as for the proof of theorem 11, we still have

*d* ≤ *ℙ*_*c,h*_(*c* has duplicate *m*-mers) + ℙ_*c,h*_(*c* is charged | no duplicate *m*-mers) and

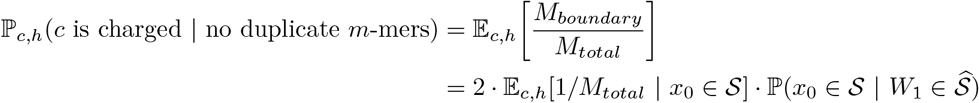

### Lemma 21

*Assuming m >* (3 + *ε*) log_*σ*_ *w*, ℙ_*c,h*_(*c has duplicate m-mers*) = *o*(1*/w*)

Proof. This proof is similar to the proof of lemma 9 from [31]. Let *i, j* ∈ *⟦*0, *w⟧* with *i < j, δ* = *j* − *i*.

If *δ < m*, 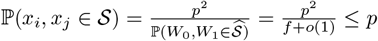

If *δ* ≥ *m*,

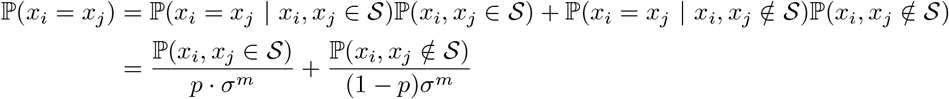

Because of lemma 17, 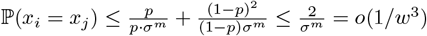 and

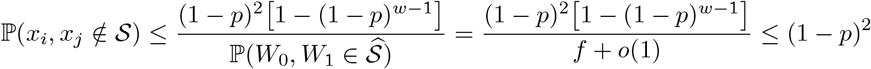

Therefore, 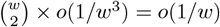

Thus ℙ_*c,h*_(*c* has duplicate *m*-mers) 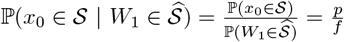

Assuming *x*_0_ is small, the *w* next *m*-mers of *c* form a *k*-mer, so we know that at least one of them is also small. Therefore,

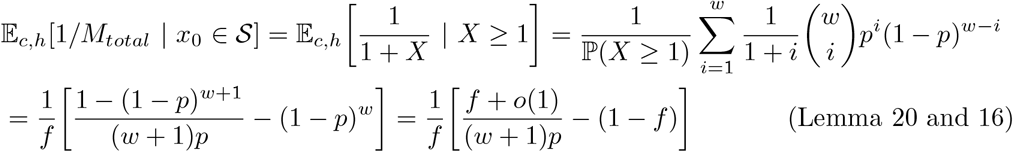

What’s more, 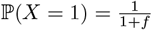. Hence,

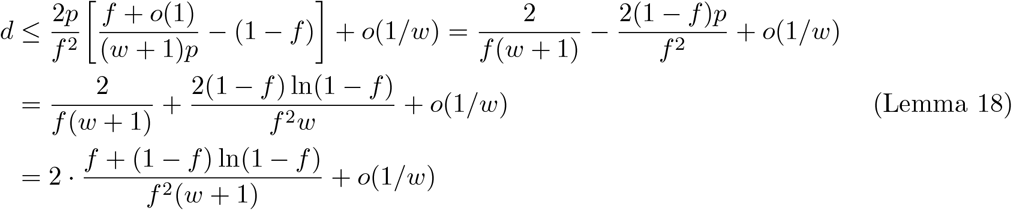

## D Proof of theorem 13

In order to compute the proportion of maximal super-*k*-mers, we adapt the proof of theorem 4 from [22].

First, we introduce a similar Markov chain representing the position *X* of the small minimizer in the *k*-mer, with an extra state ∅ when there is no small minimizer.

We reuse the following notations introduced in [22]:

*P*_*lr*_ is the proportion of left-right-max (i.e. maximal) super-*k*-mers

*P*_*l*_ is the proportion of left-max super-*k*-mers

*P*_*r*_ is the proportion of right-max super-*k*-mers

*P*_*n*_ is the proportion of non-max super-*k*-mers

By symmetry,*P*_*l*_ = *P*_*r*_, so 1− ℙ (*X* = 1)*· f ·* (1 −1*/w*) = ℙ (*X* = 1) *·* (1 + *f/w*)

What’s more, 1 + *P*_*lr*_ = *P*_*lr*_ + *P*_*l*_ + *P*_*lr*_ + *P*_*r*_ + *P*_*n*_ = 2 *· ℙ*(*X* = *w*) + *P*_*n*_ so

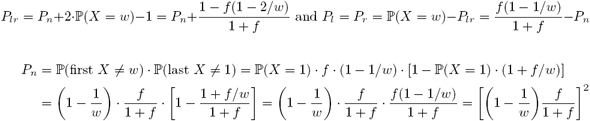

Therefore, 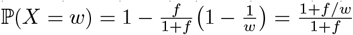 and 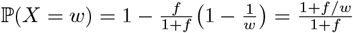

## E Proof of theorem 14

This proof generalizes the proof of theorem 11 when the minimizers are selected from a UHS

***𝒰*** with density *d****𝒰***.

First, because of independence, we have ℙ(*x*_0_ ∈ ***𝒮*** ∩ ***𝒰***) = ℙ(*x*_0_ ∈ ***𝒮***) *· ℙ*(*x*_0_ ∈ ***𝒰***) and ℙ(|***𝒮*** ∩ **𝒰** ∩ *c*| = *i*) = ∑_*n*≥*i ℙ*_(|**𝒰** ∩ *c*| = *n*) ℙ(|***𝒮*** ∩ **𝒰** ∩ *c*| = *I* | |**𝒰** ∩ *c*| = *n*)

The main change of the proof lies in the bound on the expectation:

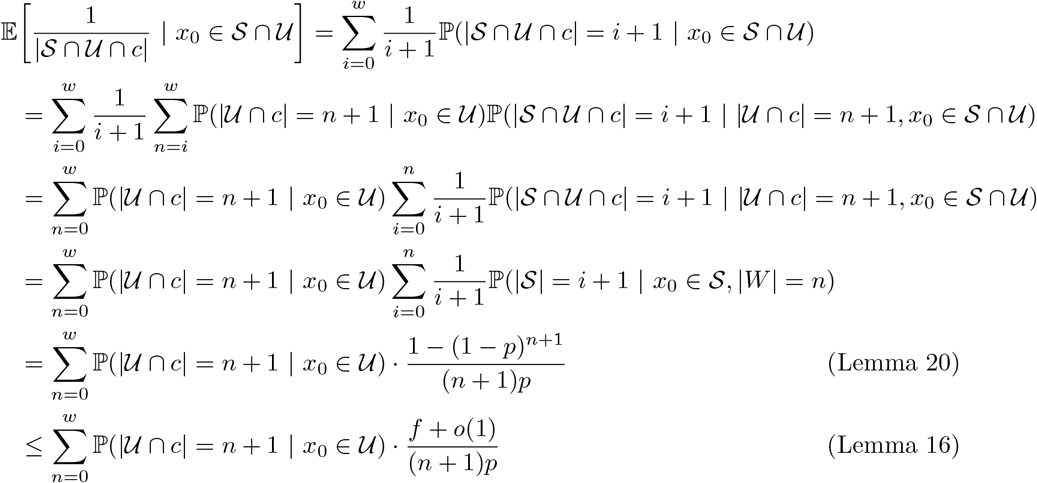

Therefore, using the same arguments as in the proof of theorem 11, we obtain

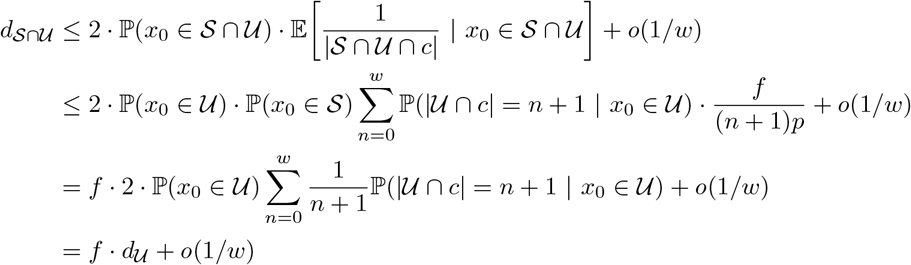

## F Additional figures and algorithms

### Algorithm 1

Comparison of two sketches

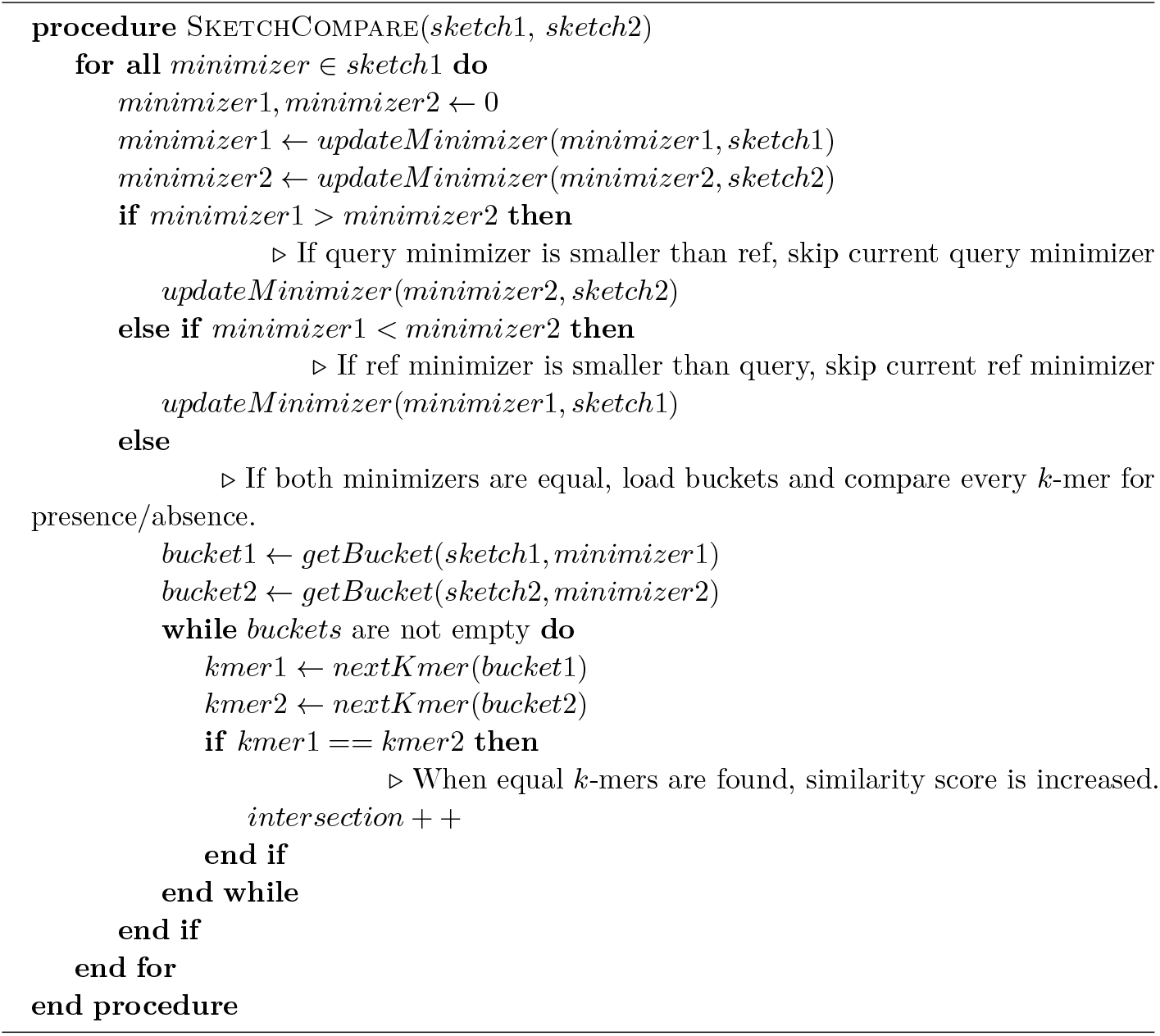

### Algorithm 2

All versus all comparison

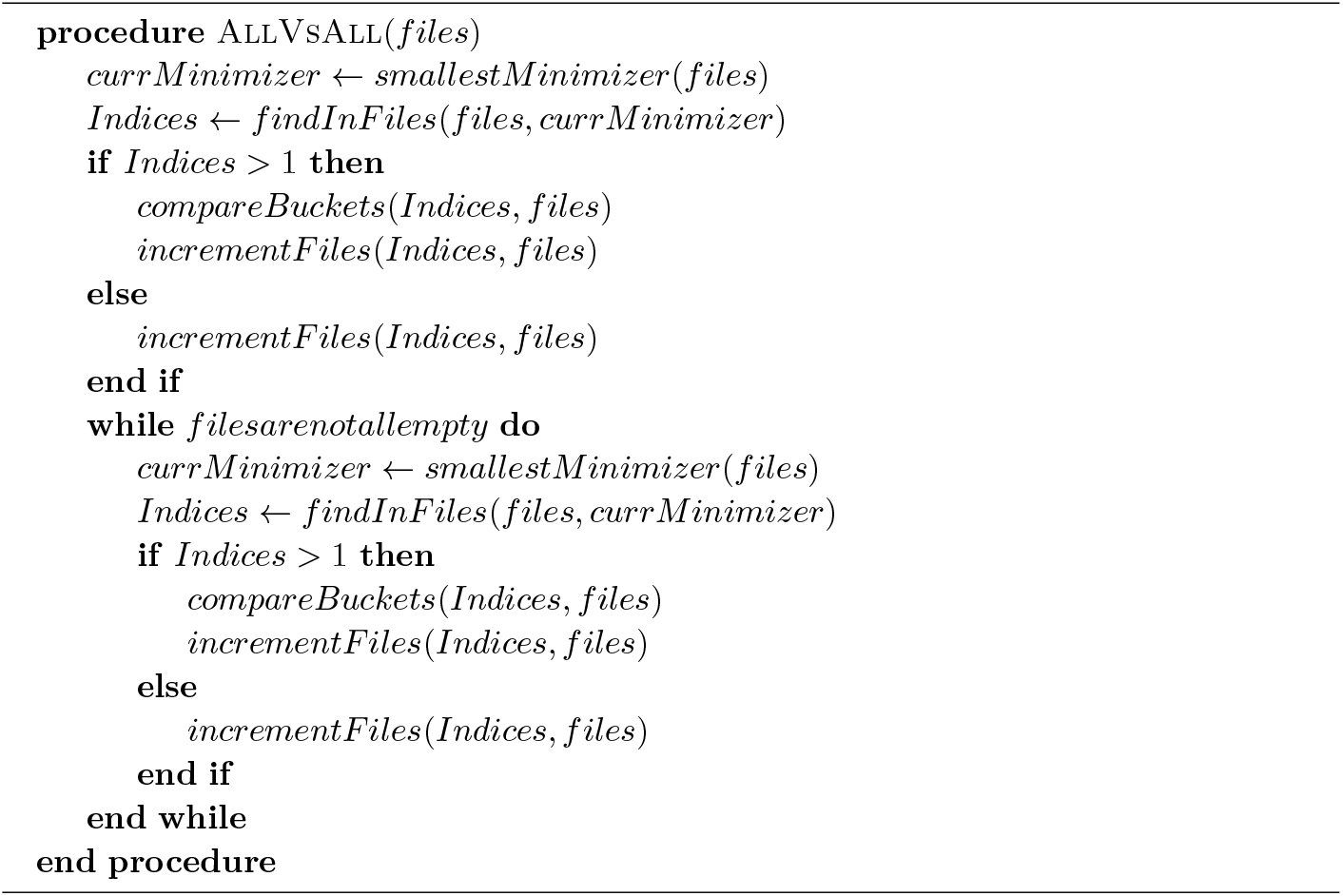

